# Membership Inference on Synthetic Single-Cell Genomic Data

**DOI:** 10.64898/2026.01.22.701160

**Authors:** Steven Golob, Patrick McKeever, Sikha Pentyala, Martine De Cock, Jonathan Peck

## Abstract

Single-cell RNA sequencing (scRNA-seq) data is subject to strict access control due to its sensitive nature, motivating the use of synthetic data generation (SDG) for privacy-preserving data sharing. We present the first adversarial privacy attack that performs meaningfully above random guessing against state-of-the-art scRNA-seq SDG methods. Our attack enables donor-level membership inference, demonstrating that leading SDG techniques fail to adequately mask which individuals were used to train the generator. We show that privacy leakage increases as the number of training donors decreases. Although the attack is designed to exploit vulnerabilities in scDesign2, we find that it also succeeds against synthetic data generated by other leading methods, including scDesign3 and scVI. This transferability indicates that an adversary can infer sensitive information from synthetic data without access to the training procedure, model parameters, or even the underlying generation algorithm. Finally, we investigate the use of perturbation with noise during the SDG process as a first-line defense, empirically evaluating its effectiveness in neutralizing the attack and its impact on utility.

## Introduction

Single-cell RNA sequencing (scRNA-seq) measures gene expression at cellular resolution (a granularity so consequential it was named Nature’s Method of the Year [29]) enabling discoveries in transcriptome profiling, personalized medicine, and tumor microenvironment characterization that bulk sequencing cannot [5, 19, 23]. Because scRNA-seq data is tied to individual donors and can reveal sensitive genetic and disease information, it is subject to strict access controls [31, 32, 33]. Indeed, even de-identified single-cell expression matrices can be linked back to individuals: one recent study re-identified donors from scRNA-seq counts via eQTL genotypes with 99.9% accuracy [43]. Because genetic variants are heritable, such exposure risks harm not only to individuals but to their biological relatives [37].

While driven by valid privacy concerns, high data access barriers create real costs. The high-friction environment for accessing genomic data stands in stark contrast to the rapid workflows that have fueled recent advances in AI. Strict data access policies induce a “privacy chill” among data custodians and constrain research progress [9]. Synthetic data generation (SDG) [17], i.e., training a generative model on real data to produce artificial samples that preserve key statistical properties such as per-gene expression distributions and inter-gene correlations [11, 47], has been proposed as a path to privacy-preserving sharing, with applications to benchmarking, rare cell-type augmentation, and data release [25, 26, 41]. Interest is not merely theoretical: the European Genome Data Infrastructure has begun releasing SDG-derived cancer genomic data to satisfy EU privacy regulations [35].

A critical question, however, is whether synthetic data actually protects donor privacy. Membership inference attacks (MIAs) that ask whether a specific individual’s data was used to train a given model are a standard auditing tool in the machine learning privacy literature [4, 38]. Yet existing MIAs on synthetic data [10, 12, 40], typically designed for demographic tabular data with dozens of features, fail to perform above random guessing on genomic data with thousands of features [34, 48]. No MIA specifically designed for synthetic genomic data has been shown to succeed. The CAMDA Health Privacy Challenge at ISBM2025^1^ highlighted this gap explicitly, calling for both private scRNA-seq SDG methods and privacy audits, and none of its participants succeeded in mounting an effective adversarial attack.

In this paper, we present an attack primarily designed around *scDesign2* and *scDesign3* (Gaussian copula variant) [41, 39], the highest-quality scRNA-seq SDG methods under the CAMDA2025 evaluation metrics [27]. These methods fit per-gene, per-cell-type marginal distributions and capture gene correlation structure via a Gaussian copula, sampling synthetic data from the learned joint distribution. We propose **scMAMA-MIA**, a single-cell– specific adaptation of the MAMA-MIA attack [10] that exploits this generative structure. Crucially, we find that it performs comparably well against all SOTA scRNA-seq SDG methods we evaluated, suggesting the vulnerability is not specific to any one model’s architecture but is a broader property of synthetic scRNA-seq data.

The remainder of this paper is structured as follows: after presenting the problem description and the proposed scMAMA-MIA attack, we demonstrate on a series of scRNA-seq datasets that scMAMA-MIA substantially outperforms existing tabular MIAs across all SDG methods tested, including scDesign2 [41], scDesign3 [39], scVI [24], and ZINB-WaVE [36]. We accompany the attack with quality evaluations of these SOTA SDG methods to characterize the privacy/utility trade-off, and an empirical study of the effect that a first-line defense of perturbing with noise during the SDG process has on said privacy/utility trade-off.

### Problem Description and Threat Models

A membership inference attack (MIA) empirically audits the privacy risk of a system ^2^ derived from real data by attempting to infer which real samples were used to train it [38]. We focus on MIAs against generative models: given synthetic dataset *D*_*synth*_ and a set of target instances *D*_*target*_, the adversary infers which targets were used to train the generator (see Fig. 1).

**Fig. 1.**
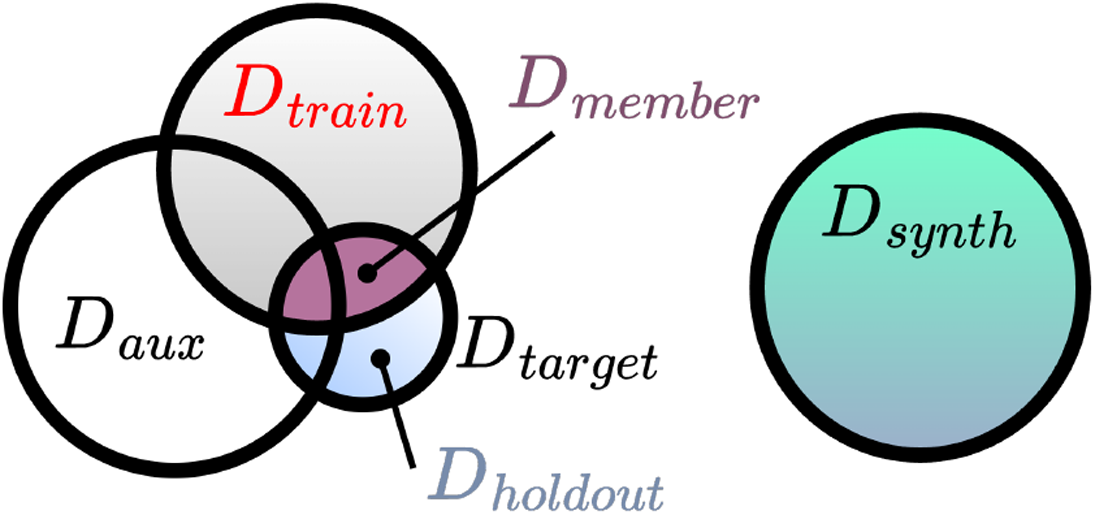
Threat model. In a MIA, the adversary has access to synthetic data *D*_*synth*_ but not training data *D*_*train*_, and must infer *D*_*train*_ ∩ *D*_*target*_ from a set of target instances *D*_*target*_. The adversary may or may not have access to auxiliary data *D*_*aux*_.

#### Synthetic Data Generation

scRNA-seq datasets have the form 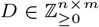, where *n* is the number of cells in the sample and *m* is the number of genes. There are |*T*| cell types and *P* donors (as in “patients”). Each row in *D* has one cell type and one donor. *D*_*t*_ denotes all the cells in *D* of type *t* while *D*_*p*_ denotes cells only from donor *p* in *D*.

A data holder with such an scRNA-seq dataset *D*_*train*_ runs an SDG algorithm, such as *scDesign2*, to create a synthetic scRNA-seq dataset *D*_*synth*_. scDesign2 [41] generates synthetic scRNA-seq counts via a two-group strategy: per-gene, per-cell-type marginals are fit via maximum likelihood over Negative Binomial, ZINB, Poisson, and Zero-Inflated Poisson families. Genes with sufficient non-zero expression (“group 2”) are jointly modeled via a Gaussian copula to preserve gene-gene correlations; the remainder (“group 1”) are sampled independently.^3^ A copula models a multivariate distribution by separating the marginal distributions of individual variables from their dependency structure [30]. Formally, given random variables *X*_1_, …, *X*_*k*_ with CDFs *F*_1_, …, *F*_*k*_, the probability integral transform (PIT) maps each *X*_*i*_*→ U*_*i*_ = *F*_*i*_(*X*_*i*_), yielding uniform marginals whose joint distribution is a copula. A Gaussian copula captures dependency via a covariance matrix *ℛ* over Gaussian-transformed marginals:

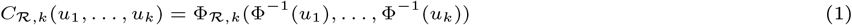

where Φ^−1^ is the inverse standard Gaussian CDF and Φ_ℛ,*k*_ is the multivariate normal CDF with covariance ℛ. To sample, one draws *Z* ∼ *𝒩* (0, *ℛ*), applies the PIT to obtain uniform *U*_*i*_ = Φ(*Z*_*i*_), then applies the inverse marginal CDFs 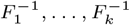 to recover samples on the original scale, preserving the correlation structure encoded in ℛ across arbitrary marginal distributions.

*scDesign3* [39] extends scDesign2 to spatial data and additional omics modalities, with a user-selectable Gaussian (default) or vine copula — the latter being substantially slower and not recommended for *>*1000 genes. scMAMA-MIA (Sec. 3) targets the Gaussian copula variant of both models; we also report performance against the vine copula variant in Sec. 4, with full adaptation left to future work.

#### Threat Models

We consider four threat models varying two axes of adversarial knowledge: whether the adversary has access to the fitted scDesign2 Gaussian copula (**white-box**, WB) or only to *D*_*synth*_ (**black-box**, BB), and whether they have access to an auxiliary real dataset from the same distribution (**+aux** / **-aux**), as in [1, 16]. The adversary observes target instances (cells) *D*_*target*_ with donor IDs and cell types, and assumes all cells from a given donor are either entirely in *D*_*train*_ or entirely held out. Although scMAMA-MIA is designed around scDesign2’s Gaussian copula, the black-box variants treat the generator as opaque; they operate on *D*_*synth*_ alone, without assuming knowledge of which SDG was used. The adversary’s task is to infer, for a given donor in *D*_*target*_, whether that donor was present in *D*_*train*_. This is critical because, if a donor’s data can be linked to the training of a synthetic cancer dataset, for example, then this would reveal that the person has cancer [48].

##### Algorithm 1

scMAMA-MIA membership inference attack, (**BB-aux**)

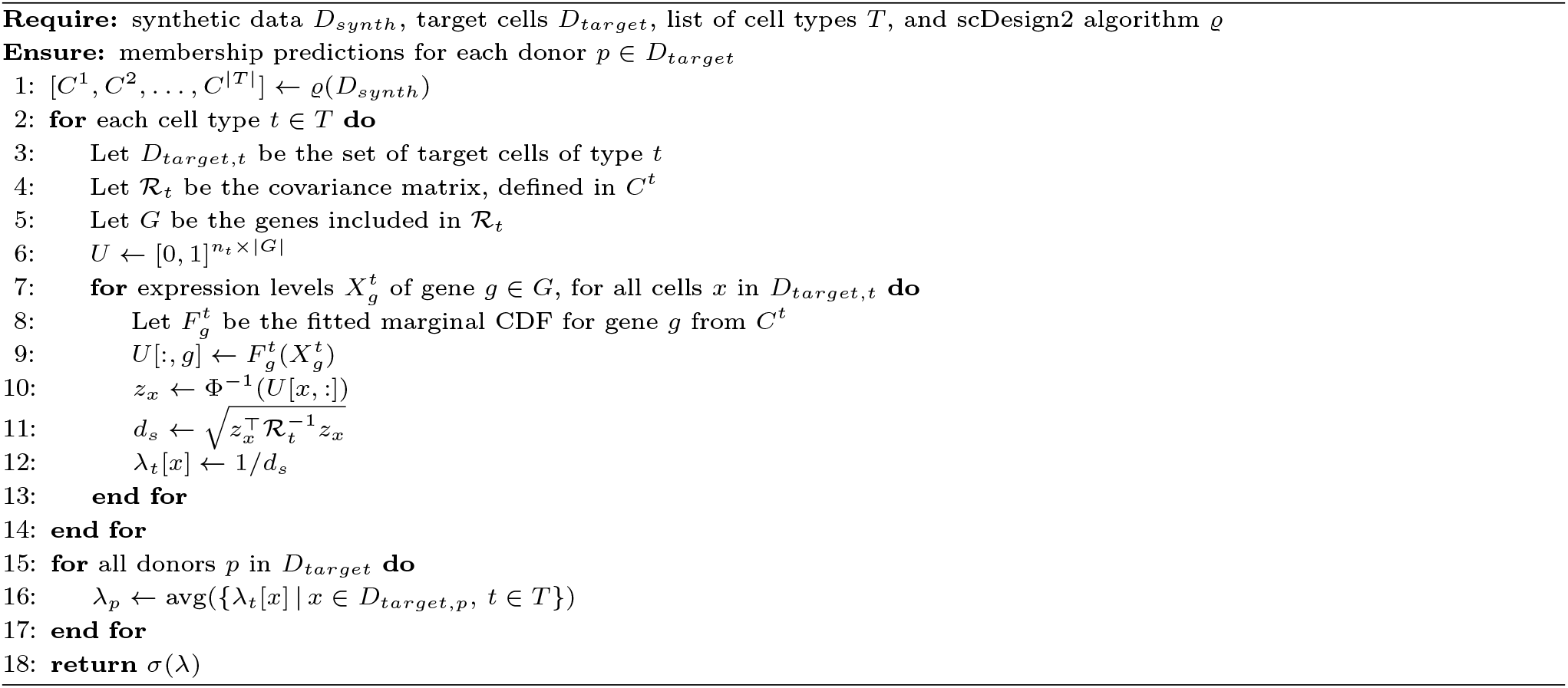

### Our Approach to Privacy Auditing: scMAMA-MIA

MAMA-MIA (Marginals Measurement Aggregation-based MIA) is a class of MIAs that exploits knowledge of an SDG’s architecture to infer membership [10]. The heuristic proceeds in three steps: (1) **identify** which statistics of the training data are preserved by the SDG (“focal-points”); (2) **simulate** the SDG to record which focal-points are selected; and (3) **aggregate** focal-point measurements on target instances to score their similarity to the synthetic data. MAMA-MIA has been shown effective against several SDGs on demographic data. We adapt the MAMA-MIA membership inference attack to scRNA-seq synthetic data, exploiting knowledge of how scDesign2 fits its generative model. We outline the three steps of the resulting scMAMA-MIA below.

#### Choice of Focal-Points

In **Step 1** of MAMA-MIA, we take the per-cell-type Gaussian copulas as focal-points: these are the direct statistical measurements taken on the real training data and preserved in the synthetic data, concentrating the synthetic data’s similarity to the real data. Each copula encodes (i) the marginal distribution parameters of “group 1” genes and (ii) a subset of these genes’ (“group 2”) pairwise covariance matrix ℛ. We use only group 2 genes in our attack. However, we also introduce an *enhanced* version that incorporates the group 1 genes’ marginals to great effect, in the Appendix.

In the *white-box* (WB) setting, the attacker directly accesses the true copula ϱ(*D*_*train*_) fit by scDesign2 on the hidden training data. In the *black-box* (BB) setting, the attacker instead constructs a proxy copula by running scDesign2 on *D*_*synth*_ and extracting the resulting copula ϱ(*D*_*synth*_) before the generation phase. This proxy is effective because scDesign2 conditions group 2 membership on gene sparsity, and its per-gene marginals include zero-inflation parameters that explicitly model this; consequently, most group 2 genes identified from the hidden training data are similarly identified from the synthetic data. **Step 2** of MAMA-MIA (simulating the SDG repeatedly to record focal-point variation) is unnecessary here, as scDesign2’s focal-point selection is deterministic and adds no privacy-related stochastic noise.

#### Measuring and Aggregating

**Step 3** is the most involved. For each cell type *t*, we measure how well a target cell’s expression levels fit the focal-point statistics extracted from the copulas. Specifically, for each group 2 gene *g* with fitted marginal CDF 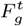 (parametrized by zero-inflation 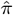, dispersion 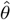, and mean 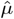), we apply the probability integral transform to map observed counts into the uniform space: 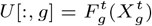, where 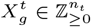 are the observed counts for gene *g* across all *n*_*t*_ target cells of type *t*. This mapping is necessary because the covariance matrix *ℛ*_*t*_ in the Gaussian copula is defined over PIT-transformed marginals, not raw counts. We then compute the Mahalanobis distance [7] between each cell’s transformed expression vector *z*_*x*_ (which is a distance from zero in the PIT-transformed space):

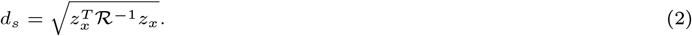

A greater *d*_*s*_ indicates that a target cell’s expression is out-of-distribution relative to the copula’s learned statistics, accounting for gene-gene covariances. We therefore take membership score 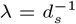 target cells close to the copula’s distribution are more likely to have contributed to it. While a high Mahalanobis distance for any individual cell may simply reflect atypical expression, *aggregating λ* over all cells from the same donor yields a stable signal. Finally, we average cell-level scores per donor and pass through a sigmoid to produce donor-level membership probabilities *p* = *σ*(*λ*) ∈ [0, 1].

#### Threat Model Variants

Algorithm 1 provides pseudocode for the **BB-aux** variant of scMAMA-MIA, i.e., assuming the least adversarial knowledge. If more knowledge is available to the adversary, in the form of either the trained SDG model and/or an additional auxiliary dataset, then the scMAMA-MIA attack is further strengthened as follows:

- **WB vs. BB**: In the WB setting, the adversary has access to the trained SDG model, and so ϱ(*D*_*train*_) is used directly instead of the synthetic copula; in the BB setting, it is replaced by the proxy ϱ(*D*_*synth*_). Note that the **WB** variant of scMAMA-MIA is not applicable to other SDG methods, since they do not produce ϱ(*D*_*train*_).
- *±***aux**: Without auxiliary data, the membership score is 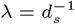. When an auxiliary real dataset *D*_*aux*_ is available, the adversary additionally computes *d*_*a*_ (i.e. the Mahalanobis distance under ϱ(*D*_*aux*_)) and sets *λ* = *d*_*a*_*/*(*d*_*s*_ + *d*_*a*_). This contrasts how well a cell fits the synthetic copula against a held-out real copula, substantially improving attack accuracy in practice. Also, for the
- +**aux** variant, the gene set *G* becomes the intersection of “group 2” genes present in *both* copulas; on average, a large portion (86.6%) of group 2 genes appear in both.

### Experiments

We evaluate scMAMA-MIA on synthetic data from three real scRNA-seq datasets across six SDG methods, averaging all results over five trials per configuration (each with a different donor sample for training, holdout, and auxiliary sets^4^). Code is available at https://github.com/steveng9/scRNA-seq_privacy_audits. ^5^

#### Datasets, SDG Methods, and Experimental Setup

We evaluate on three scRNA-seq datasets of varying scale, demographic composition, and donor count (Table 1). **OneK1K** [46] contains peripheral blood mononuclear cells from 981 donors across 14 cell types, and served as the primary benchmark in the CAMDA2025 Health Privacy Challenge (Track 2). **AIDA** [20] comprises healthy immune cells from 508 donors across 33 cell types and seven Asian population groups from five countries; its demographic breadth heightens its sensitivity. **HFRA** [22] is a Human Fetal Retina Atlas from just 22 donors across 9 cell types, yielding substantially more cells per donor than the other two. Both AIDA and HFRA are retrieved from the CZ Cell x Gene database.^6^

**Table 1.** Dataset statistics and scDesign2 gene group assignments. Group 1 genes are modeled via independent marginals; group 2 genes are jointly modeled via the Gaussian copulas. HVGs are the highly variable genes retained for all experiments.

| Dataset | Donors | Cells | Cell types | HVGs | scDesign2 genes (mean ± std) | |
| --- | --- | --- | --- | --- | --- | --- |
|  |  |  |  |  | group 1 | group 2 |
| OneK1K | 981 | 1,267,733 | 14 | 1,118 | 864 ± 284 | 145 ± 73 |
| AIDA | 508 | 1,058,909 | 33 | 2,021 | 1,214 ± 465 | 446 ± 98 |
| HFRA | 22 | 108,838 | 9 | 3,664 | 3,044 ± 816 | 363 ± 110 |

We follow [41] and select highly variable genes (HVGs) via `sc.pp.highly_variable_genes` in Scanpy with `min_mean`=0.0125, `max_mean`=3, and `min_disp`=0.5. UMAPs [14] of representative training and synthetic samples are provided in Figure 3.

In addition to **scDesign2** and **scDesign3** (Gaussian and vine copula variants; Sec. 2), we evaluate against four further SOTA SDG methods: **scVI** [24], a variational autoencoder with a ZINB observation model; **scDiffusion** [25], a diffusion model conditioned on cell-type embeddings; **ZINB-WaVE** [36], a zero-inflated factor model; and **NMF** [44], which decomposes expression data via non-negative matrix factorization and samples synthetic cells from per-cluster distributions.

We experiment with training sets of {2, 5, 10, 20, 50, 100, 200} donors, retaining all cells per sampled donor. The target set *D*_*target*_ is a disjoint same-size sample concatenated with the training data, so that exactly half of target donors are members—the standard MIA setup [1, 16, 28, 45]. Auxiliary data *D*_*aux*_ is sampled similarly (minimum 10 donors) and need not be disjoint from train or holdout. HFRA experiments are limited to ≤11 donors given its small donor pool.

#### Success Rate of scMAMA-MIA against scDesign2

Figure 2 shows ROC AUC scores for donor-level membership inference attacks against scDesign2-generated synthetic data, across all three datasets and varying donor sample sizes. We compare scMAMA-MIA (black-box, with auxiliary data) against five attacks from the CAMDA2025 benchmark: LOGAN [13], GAN-Leaks (calibrated) [6], DOMIAS with KDE [42], and Monte Carlo [15]. All attacks operate under the black-box threat model with access to an auxiliary dataset.

**Fig. 2.**
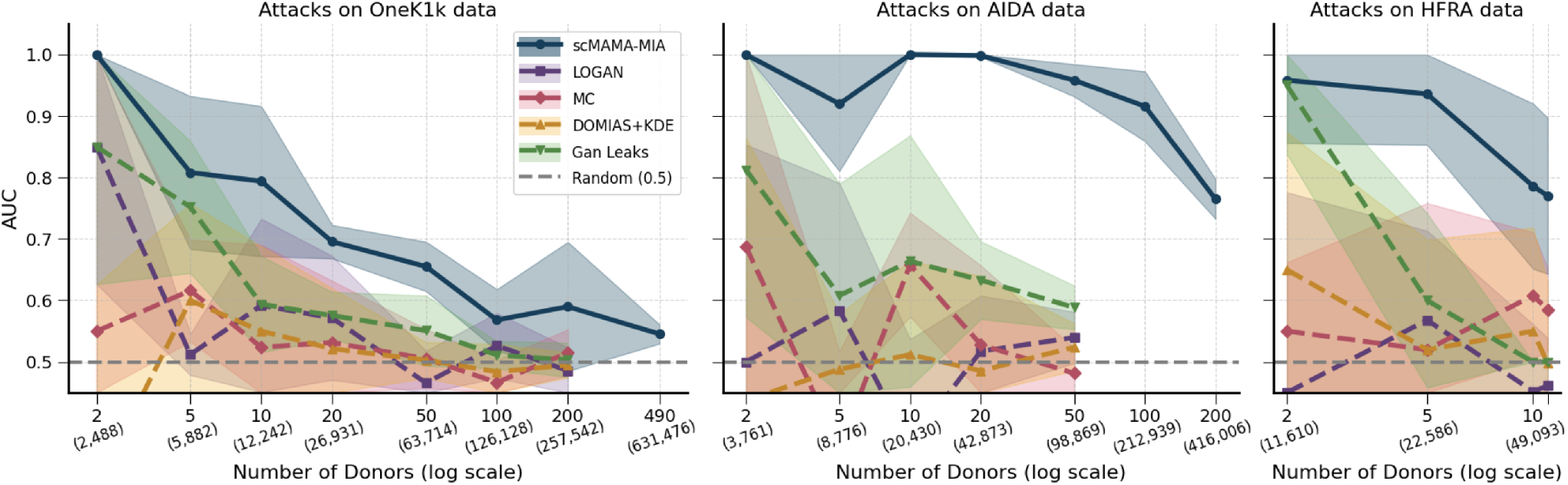
Success rate of scMAMA-MIA and baseline attacks, measured in AUC on **scDesign2** generated synthetic datasets for OneK1K, AIDA, HFRA, and for a varying number of donors. (The average number of cells for samples is given in parenthesis.) All attacks are in the black-box (BB) threat model with access to an auxiliary dataset. Color bands denote *±*1 standard deviation from the mean AUC.

**Fig. 3.**
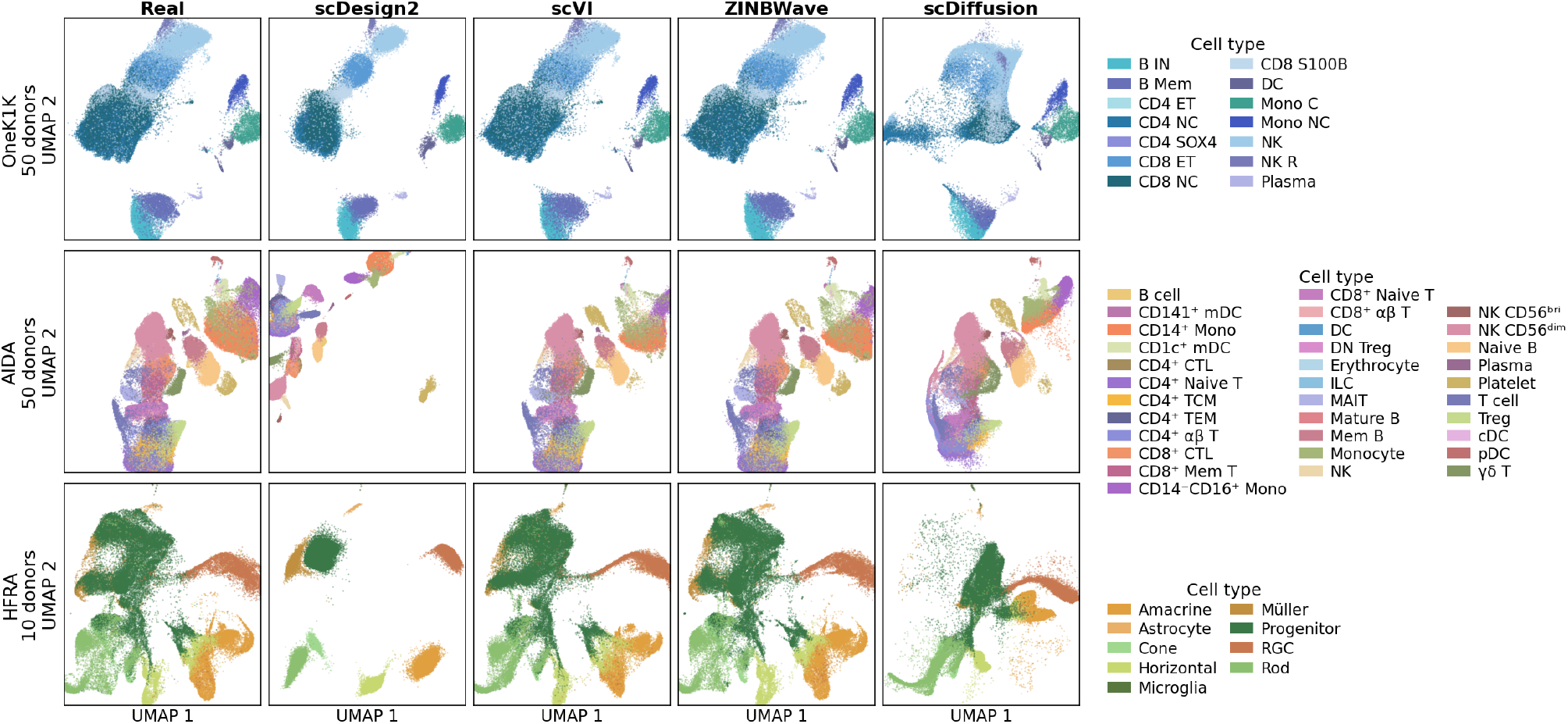
UMAPs for real data samples and the corresponding synthetic data (computed jointly) to illustrate how much information each SDG method is able to preserve.

scMAMA-MIA substantially outperforms all baselines across every donor count and dataset. Among the baselines, GAN-Leaks (calibrated, i.e., also leverages *D*_*aux*_) tends to be the strongest competitor, yet still falls well short of scMAMA-MIA, particularly at larger donor sizes, where most baselines regress toward random-chance performance. Notably, merely obtaining meaningful attack AUCs at scale with the baseline MIAs required considerable engineering effort: we implemented batched versions of GAN-Leaks, DOMIAS+KDE, and Monte Carlo to handle larger cohorts; even so, these attacks could not complete ^7^ on AIDA above 50 donors or OneK1K above 200 donors, which accounts for the gaps in Figure 2. In terms of computational efficiency, scMAMA-MIA is consistently faster than all baselines across donor sizes, with speedups ranging from 2.6× to 12.3× at 50 donors, while LOGAN remains the only faster method (≈2–3×) owing to its substantially lighter inference pipeline.

A particularly striking result is that scMAMA-MIA achieves perfect or near-perfect AUC (1.00) at the smallest donor counts across all three datasets. On AIDA, this extends to datasets with 10 and 20 donors (roughly 43,000 cells) as well. This is notable precisely because the black-box-with-auxiliary-data setting (BB+aux) is only the third most powerful threat model we evaluate; achieving perfect inference under this setting reflects how the signal sharpens in scDesign2 when donor cohorts are small. The success of scMAMA-MIA in general can be attributed to its exploitation of the specific way scDesign2 fits a model on real data: a structural vulnerability that the existing attacks, which treat the generator as a generic black box, are entirely unable to leverage.

Figure 4 examines how scMAMA-MIA performs against scDesign2 across four threat models on the OneK1K dataset: black-box with auxiliary data (BB+aux), black-box without auxiliary data (BB−aux), white-box with auxiliary data (WB+aux), and white-box without auxiliary data (WB−aux). As expected, WB+aux is the strongest overall, and auxiliary data generally improves attack success. However, the most practically interesting comparison is between WB−aux and BB+aux: across donor sizes, WB−aux consistently outperforms BB+aux, despite having access to *no* auxiliary population data. This suggests that white-box access to the generator’s learned parameters is more informative than an auxiliary dataset: an attacker who can inspect the model directly gains more than one who cannot, even if the latter has supplementary reference data at their disposal.

**Fig. 4.**
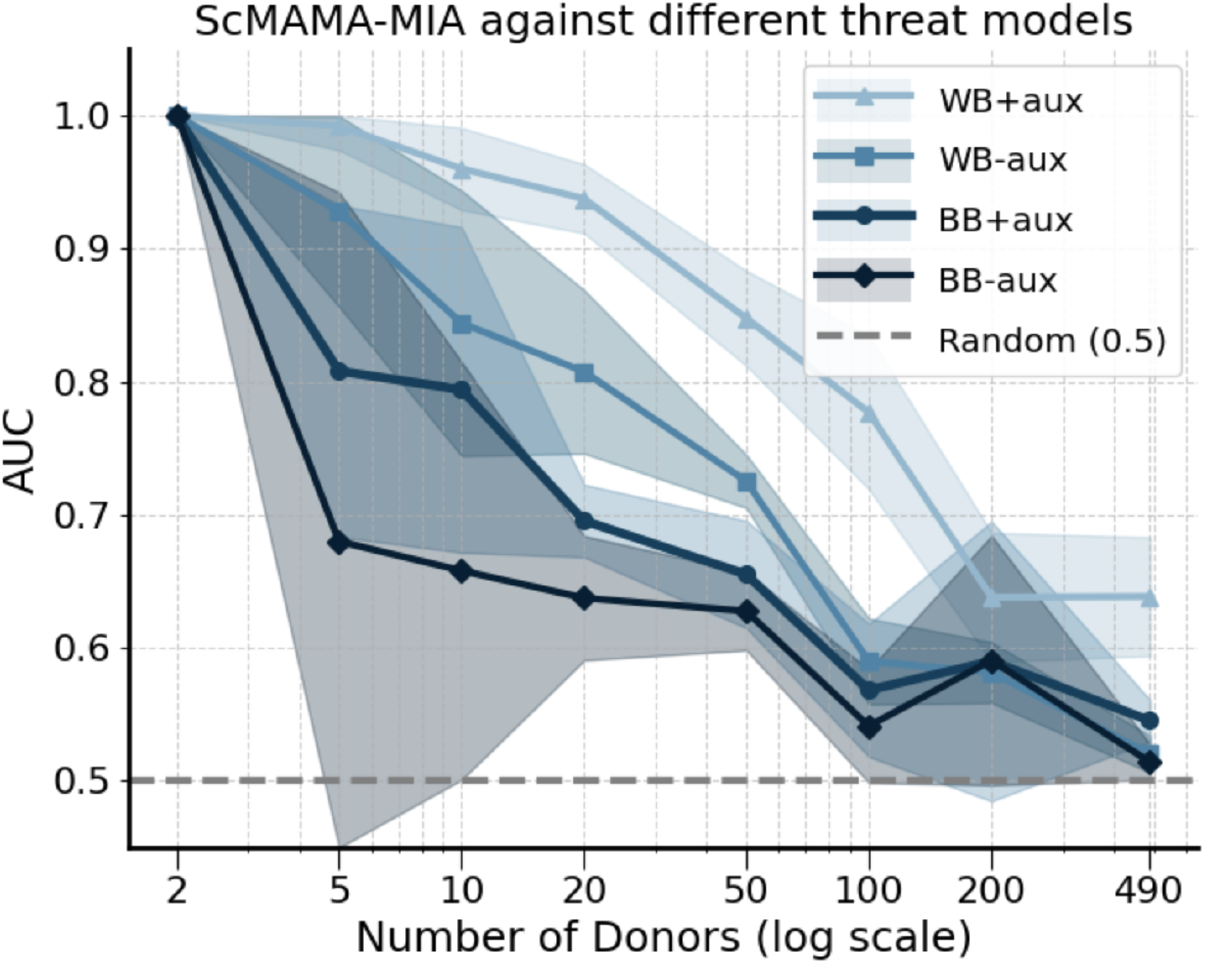
Success rate of scMAMA-MIA under different threat models, including white-box (WB) and black-box (BB) threat models, with and without auxiliary data. Conducted on **scDesign2** generated synthetic data for OneK1K.

Attack AUCs on HFRA are somewhat lower than on the other two datasets, peaking at 0.95 for the strongest threat models and declining more quickly with donor count. We observe in Section 4.3 that scDesign2’s synthetic data quality is also lower for HFRA, which likely accounts for this discrepancy; the weaker learned representation provides a less reliable signal for membership inference.

We validate scMAMA-MIA via the Mahalanobis distance distributions underlying the attack (step 3). Figure 5 shows ridge distributions of these distances for member and non-member cells, per cell type and dataset. These distances quantify how far a cell’s expression profile lies from the copula covariance structure learned from training data. Member cells consistently have lower mean distances than non-members, reflecting their contribution to the fitted copula, though the distributions overlap at the single-cell level. This overlap motivates aggregating evidence across many cells from a donor, enabling reliable membership inference via pooling. Distances are drawn from one random trial (50 donors for OneK1K, 20 for AIDA, 10 for HFRA). Variation in distance magnitude across cell types is an open question and does not affect our conclusions.

**Fig. 5.**
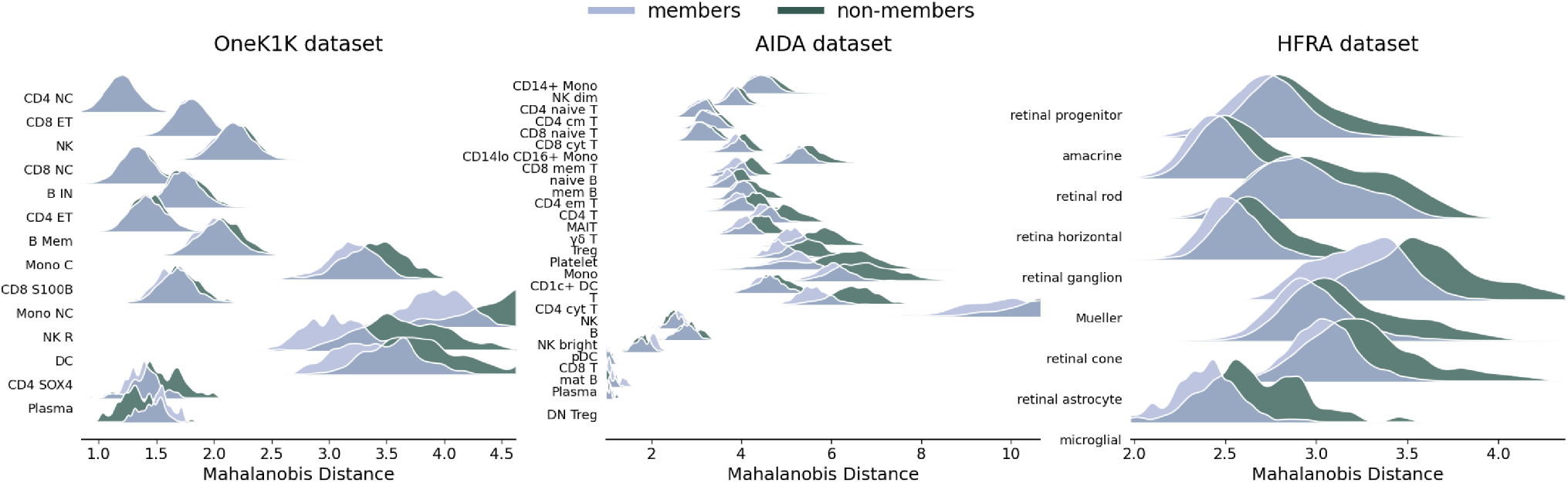
The distribution of Mahalanobis distances taken during an attack on **scDesign2**, between member and non-member cells and the copula’s covariance matrix ℛ learned from the training data. The trend of member distances being less than non-member distances validates why our attack is effective.

In the appendix, Section A, we provide attack results across demographic subgroups, sex and ethnicity, to show how privacy is unequally distributed across population groups in synthetic data generation.

#### Success Rate of scMAMA-MIA against other SDG Methods

Having established scMAMA-MIA’s effectiveness against scDesign2, we evaluate whether this behavior generalizes across other SDG methods. Table 2 shows that it does: across all five high-fidelity SDGs (scDesign2, scDesign3-G, scDesign3-V, scVI, and ZINB-WaVE), BB+aux AUC ranges from 0.59 to 0.66, well above chance. An enhanced variant incorporating auxiliary per-gene likelihood evidence (Appendix, Algorithm 2) further increases AUC up to 0.79, indicating that this signal is particularly effective against copula-based models.

**Table 2.** Fidelity and privacy for all non-DP SDG methods (OneK1K, 50 donors). BB*±*aux: scMAMA-MIA black-box attack without/with auxiliary data; CB: *Enhanced* (uses BB+aux threat) variant adding per-gene log-likelihood ratio evidence (see appendix).

| SDG Method | Fidelity |  |  | Privacy (scMAMA-MIA AUC) |  |  |
| --- | --- | --- | --- | --- | --- | --- |
| | LISI↑ | ARI↑ | MMD$\times 10^3 \downarrow$ | BB-aux | BB+aux | *enhanced* |
| scDesign2 | 0.87 ± .00 | 0.48 ± .02 | 0.45 ± .03 | 0.63 ± .03 | 0.66 ± .04 | 0.79 ± .06 |
| scDesign3-G | 0.88 ± .00 | 0.46 ± .04 | 0.28 ± .03 | 0.61 ± .03 | 0.65 ± .03 | 0.78 ± .07 |
| scDesign3-V | 0.88 ± .00 | 0.46 ± .03 | 0.31 ± .01 | 0.61 ± .02 | 0.64 ± .04 | 0.79 ± .06 |
| scVI | 0.90 ± .00 | 0.40 ± .05 | 0.28 ± .03 | 0.57 ± .03 | 0.59 ± .05 | 0.69 ± .09 |
| ZINB-WaVE | 0.89 ± .01 | 0.45 ± .03 | 1.02 ± 1.66 | 0.59 ± .04 | 0.66 ± .08 | 0.78 ± .12 |
| NMF | 0.18 ± .04 | 0.29 ± .02 | 27.56 ± 2.99 | 0.55 ± .05 | 0.56 ± .05 | 0.51 ± .08 |
| scDiffusion | 0.49 ± .01 | 0.39 ± .05 | 45.69 ± 3.99 | 0.48 ± .05 | 0.47 ± .03 | 0.47 ± .02 |

The two exceptions, NMF and scDiffusion, yield AUCs near 0.5 across all variants. However, this does not indicate privacy protection: both methods exhibit substantially degraded fidelity (NMF: LISI = 0.18, MMD = 27.56; scDiffusion: LISI = 0.49, MMD = 45.69), implying that membership inference fails primarily because the synthetic data is weakly informative about its training distribution.

This highlights a key interpretive principle: low attack AUC is only meaningful as a privacy signal when accompanied by high-fidelity synthetic data. We quantify fidelity using LISI (neighborhood mixing between real and synthetic cells; higher is better), ARI (cluster agreement; higher is better), and MMD (distributional distance; lower is better) [21, 18]. Under these metrics, the copula- and VAE-based methods (scDesign2, scDesign3-G, scDesign3-V, scVI, ZINB-WaVE) consistently occupy the high-fidelity regime (LISI ≥ 0.87, MMD ≤ 1.02) while remaining vulnerable to scMAMA-MIA. Among them, scVI exhibits the lowest attack AUC (BB+aux= 0.59), suggesting modest robustness from its latent variational structure but no effective immunity.

Across datasets, deviations in attack behavior are explained by fidelity differences: HFRA exhibits lower AUCs consistent with reduced synthetic data quality (Table 5), and its UMAP structure (Figure 3) shows weak resolution of fine-grained retinal subtypes, jointly degrading fidelity and attack signal. Interestingly, AIDA UMAPs reveal low global real–synthetic overlap for scDesign2, yet scMAMA-MIA still achieves high AUC, indicating that low-dimensional visual similarity is not predictive of membership leakage. Instead, the Gaussian copula covariance structure appears to preserve donor-specific statistical fingerprints even when global embeddings suggest distributional mismatch.

#### Effect of Noise on Data Quality and Defending Privacy

We investigate stochastic perturbation of the scDesign2 covariance estimation as a first-line privacy defense, empirically characterizing the resulting privacy–utility tradeoff. For each cell type’s Gaussian copula covariance matrix, we inject symmetric Gaussian noise: each entry *ℛ*_*jk*_ receives additive noise drawn from *𝒩* (0, *σ*^2^), where

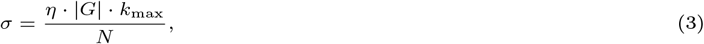

with |*G*| the number of copula genes, *k*_max_ the maximum number of cells contributed by any single donor to this cell type, and *N* the total training cells for this cell type. The scaling by |*G*| reflects a conservative, Frobenius-norm-based calibration: rather than protecting each covariance entry individually, *σ* is set proportional to the matrix displacement when one donor is removed, which aggregates over all |*G*|^2^ entries and yields a per-entry noise level |*G*| times larger than a purely entry-wise bound. The noised matrix *E* is symmetrized (i.e. (*E* + *E*^*T*^)*/*2), projected to the nearest positive semidefinite matrix, and re-normalized to a correlation matrix. We sweep *η* ∈ {10^−4^, …, 10^1^}, spanning negligible noise to well beyond typical gene–gene correlations of 0.05–0.30.

Table 3 shows the results on OneK1K at 50 donors. The tradeoff is gradual and well-behaved: fidelity (LISI, ARI, MMD) remains essentially unchanged through *η* = 10^−2^, then degrades meaningfully at *η* = 10^−1^ and collapses by *η* = 1. Attack AUC mirrors this pattern, declining from 0.66 at baseline to 0.54 (near chance) at *η* = 10^−1^, with no further gain at higher noise. This identifies *η* ≈ 10^−1^ as a practical crossover: the last setting at which meaningful privacy improvement is achievable before fidelity degrades substantially.

**Table 3.** Privacy–utility tradeoff for scDesign2 (OneK1K data, 50 donors). *η* Gaussian noise-to-signal parameter added to the copula covariance (larger = more noise). LISI ↑, ARI ↑, MMD ↓: fidelity metrics (see Section 4.3).

| Privacy Param | LISI↑ | ARI↑ | MMD$\times 10^3 \downarrow$ | BB-aux | BB+aux |
| --- | --- | --- | --- | --- | --- |
| $\eta = 0$ (least privacy) | $0.87 \pm .00$ | $0.48 \pm .02$ | $0.45 \pm .03$ | $0.63 \pm .03$ | $0.66 \pm .04$ |
| $\eta = 10^{-4}$ | $0.87 \pm .00$ | $0.49 \pm .01$ | $0.44 \pm .02$ | $0.63 \pm .03$ | $0.64 \pm .04$ |
| $\eta = 10^{-3}$ | $0.87 \pm .00$ | $0.47 \pm .04$ | $0.44 \pm .04$ | $0.62 \pm .03$ | $0.63 \pm .04$ |
| $\eta = 10^{-2}$ | $0.85 \pm .00$ | $0.46 \pm .04$ | $0.42 \pm .02$ | $0.59 \pm .02$ | $0.60 \pm .03$ |
| $\eta = 10^{-1}$ | $0.72 \pm .01$ | $0.51 \pm .03$ | $0.47 \pm .02$ | $0.55 \pm .03$ | $0.54 \pm .01$ |
| $\eta = 1$ | $0.62 \pm .01$ | $0.47 \pm .03$ | $0.53 \pm .03$ | $0.52 \pm .02$ | $0.49 \pm .03$ |
| $\eta = 10^1$ | $0.61 \pm .01$ | $0.54 \pm .02$ | $0.55 \pm .01$ | $0.52 \pm .02$ | $0.49 \pm .02$ |

Extending this approach to formal donor-level differential privacy [8] faces fundamental obstacles: the sheer number of parameters to be protected—the |*G*| *×* |*G*| covariance entries plus hundreds of per-gene marginal parameters, across cell types where individual donors may contribute thousands of cells—makes calibrating DP noise to a reasonable privacy budget without catastrophic utility loss an open problem. We leave a rigorous DP mechanism for single-cell SDGs to future work.

## Discussion

We introduced scMAMA-MIA, the first membership inference attack effective against synthetic scRNA-seq data, exploiting the donor-specific statistical fingerprints embedded in scDesign2’s Gaussian copula. Against all three datasets, scMAMA-MIA substantially outperforms existing MIA baselines under all threat models, achieving perfect AUC at small donor counts even under black-box conditions. It generalizes to other SDG methods despite being designed around the copula structure, suggesting that the underlying vulnerability is broader than any single generative model.

Our quality analysis clarifies an important interpretive point: low attack AUC is only a meaningful privacy signal when the synthetic data is also high-fidelity. Methods such as NMF and scDiffusion resist the attack primarily because their synthetic data is too dissimilar from the real data to carry a membership signal, not because they offer genuine privacy protection. Among high-fidelity SDGs, scVI shows the most modest vulnerability, warranting further investigation.

Our noise injection experiments demonstrate a gradual, well-behaved privacy–utility tradeoff for scDesign2: fidelity intact at low noise levels and degrades as noise increases, with attack AUC approaching chance near *η* ≈ 10^−1^ before fidelity collapses. This is an encouraging result for practitioners seeking defenses. However, extending this to formal donor-level differential privacy remains an open problem, given the scale of parameters involved, and we leave it for future work.

Practically, we recommend that model parameters (e.g. copula covariances and marginal fits) never be released alongside synthetic data, as white-box access consistently amplifies attack success. Increasing the donor cohort size also reduces leakage, as individual expression profiles are increasingly able to “hide among the crowd.” We hope this work motivates a more rigorous treatment of privacy in synthetic genomic data and raises the bar for what constitutes an adequate privacy guarantee.

### Limitations

Despite these findings, scMAMA-MIA is primarily designed for covariance-based SDGs (e.g. scDesign2 / scDesign3-G), and while it transfers empirically to other models, its behavior is not theoretically characterized across diffusion-or likelihood-free generators. We also focus on a fixed donor-level threat model and do not consider more adaptive adversaries or richer external genomic reference integration.

## Author contributions statement

All authors worked together on the overall design of the solution. Steven Golob implemented the attack, designed and ran the experiments. All authors discussed results and wrote the manuscript together.

## Acknowledgments

Steven Golob is supported by an NSF CSGrad4US fellowship and an IEEE Computational Intelligence Society Graduate Student Research Grant. This material is based upon work supported by the National Science Foundation under Grant No. 2451163 and No. 2523406, and by NSF NAIRR 240485 (Cloudbank AWS) and NSF NAIRR 240091 (TACC Frontera). This research was, in part, funded by the National Institutes of Health (NIH) Agreement No. 1OT2OD032581. The views and conclusions contained in this document are those of the authors and should not be interpreted as representing the official policies, either expressed or implied, of the NIH.

## Algorithm 2

scMAMA-MIA (**BB-aux**), ***enhanced*** with copula’s one-way marginal parameters

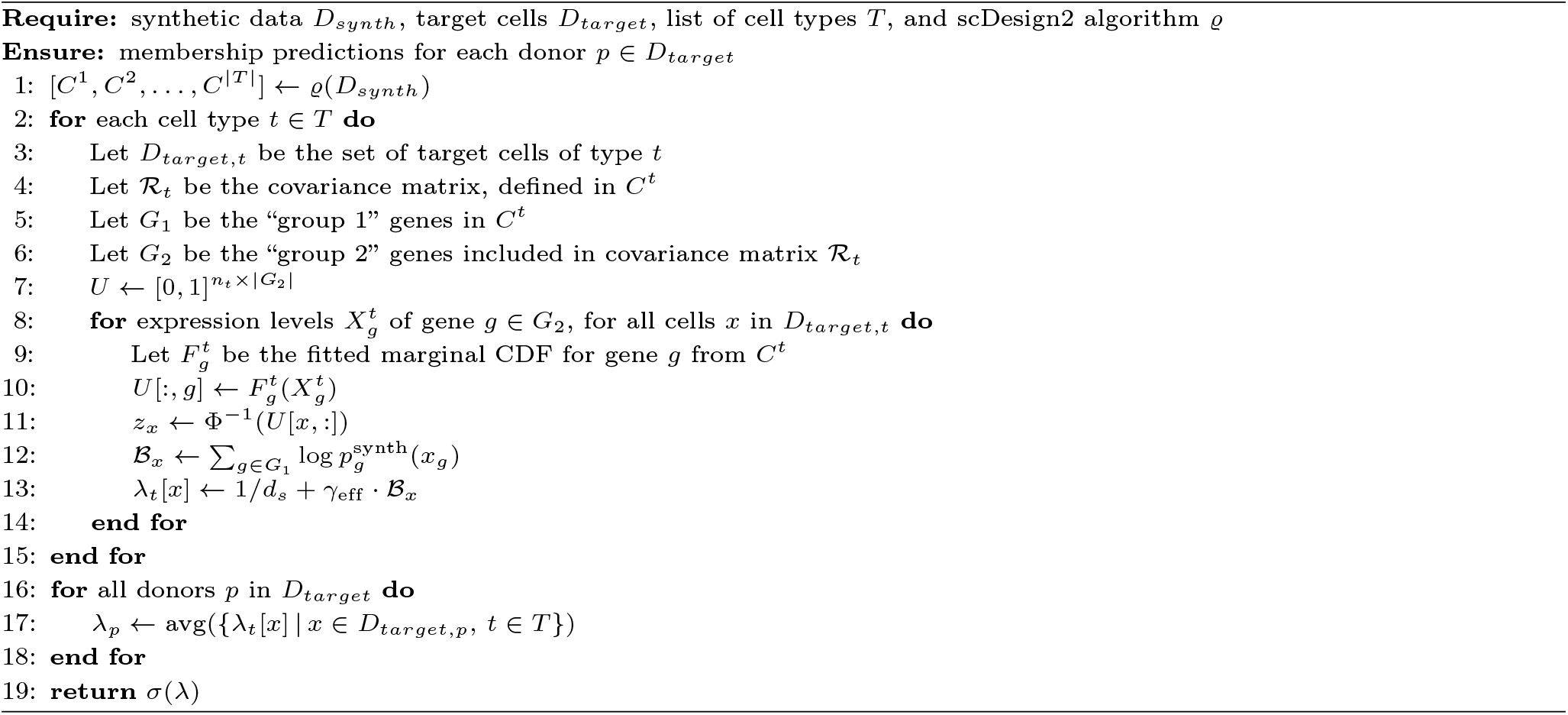

## By Demographic Subgroup

Privacy leakage may be more severe for demographic subgroups underrepresented in training data [3]. Table 4 shows donor-level AUC broken down by sex and ethnicity on the AIDA dataset (BB+aux, 20 donors). For scDesign2 and scDesign3-G, scMAMA-MIA achieves perfect AUC uniformly across all subgroups. Neural network- and matrix-factorization-based SDGs exhibit substantially higher variance in attack success across subgroups — likely because these models are more sensitive to distributional shifts between groups, amplifying differences in data quality and representation into unequal privacy exposure. For sex, privacy leakage tends to be higher for females (55.1% of donors) than for males — most clearly in scVI (0.74 vs. 0.59) and ZINBWave (0.84 vs. 0.68) — consistent with the hypothesis that overrepresented groups are more vulnerable due to better model fit. For ethnicity, AUCs fluctuate considerably across groups and SDGs (e.g. scDiffusion ranges from 0.17 for Singaporean Indian to 0.80 for Singaporean Chinese), with no consistent relationship to group size; cell-type composition and batch effects are likely confounders.

**Table 4.** scMAMA-MIA (BB+aux) disparate-impact breakdown by sex and ethnicity on the aida dataset, averaged over 5 samples with 20 donors. Subgroup cell-count prevalence in parentheses.

| Subgroup | scDesign2 | scDesign3-G | scDesign3-V | scVI | scDiffusion | NMF | ZINBWave |
| --- | --- | --- | --- | --- | --- | --- | --- |
| *Sex* |  |  |  |  |  |  |  |
| female (55.1%) | 1.00±0.00 | 1.00±0.00 | 0.66±0.32 | 0.74±0.13 | 0.55±0.24 | 0.68±0.20 | 0.84±0.10 |
| male (44.9%) | 1.00±0.00 | 1.00±0.00 | 0.63±0.29 | 0.59±0.12 | 0.48±0.14 | 0.54±0.12 | 0.68±0.07 |
| *Ethnicity* |  |  |  |  |  |  |  |
| Japanese (27.1%) | 1.00±0.00 | 0.99±0.01 | 0.57±0.34 | 0.65±0.18 | 0.63±0.20 | 0.64±0.19 | 0.73±0.16 |
| Korean (34.6%) | 1.00±0.00 | 1.00±0.00 | 0.70±0.32 | 0.69±0.15 | 0.35±0.15 | 0.62±0.19 | 0.84±0.09 |
| Singaporean Chinese (11.7%) | 1.00±0.00 | 1.00±0.00 | 0.85±0.20 | 0.69±0.11 | 0.80±0.19 | 0.60±0.24 | 0.85±0.18 |
| Singaporean Indian (10.1%) | 1.00±0.00 | 1.00±0.00 | 0.50±0.41 | 0.67±0.24 | 0.17±0.24 | 0.71±0.16 | 0.71±0.26 |
| Singaporean Malay (9.5%) | 1.00±0.00 | 1.00±0.00 | 0.58±0.43 | 0.80±0.40 | 0.32±0.19 | 0.53±0.45 | 1.00±0.00 |

**Table 5.** Quality metrics (LISI, ARI, and MMD) for **scDesign2** across datasets and donor counts. Values are mean *±* std over 5 trials. ↑ = higher is better; ↓ = lower is better. MMD values scaled by 10^3^. HFRA donor pool is limited to 22 donors.

| Donors | OneK1K |  |  | AIDA |  |  | HFRA |  |  |
| --- | --- | --- | --- | --- | --- | --- | --- | --- | --- |
| | LISI ↑ | ARI ↑ | MMD $\times 10^3 \downarrow$ | LISI ↑ | ARI ↑ | MMD $\times 10^3 \downarrow$ | LISI ↑ | ARI ↑ | MMD $\times 10^3 \downarrow$ |
| 2 | 0.91±.01 | 0.54±.16 | 0.03±.00 | 0.87±.01 | 0.76±.08 | 0.41±.05 | 0.56±.02 | 0.41±.14 | 0.51±.11 |
| 5 | 0.90±.01 | 0.53±.04 | 0.03±.01 | 0.83±.02 | 0.71±.11 | 0.35±.06 | 0.37±.07 | 0.44±.08 | 0.59±.11 |
| 10 | 0.88±.01 | 0.50±.06 | 0.04±.01 | 0.81±.01 | 0.74±.06 | 0.38±.04 | 0.27±.04 | 0.48±.06 | 0.65±.05 |
| 20 | 0.88±.00 | 0.50±.04 | 0.04±.00 | 0.78±.02 | 0.74±.04 | 0.35±.04 | 0.31±.06 | 0.45±.07 | 0.63±.12 |
| 50 | 0.87±.00 | 0.48±.02 | 0.04±.00 | 0.75±.01 | 0.77±.02 | 0.35±.02 | — | — | — |
| 100 | 0.86±.00 | 0.48±.02 | 0.04±.00 | 0.73±.00 | 0.74±.05 | 0.34±.02 | — | — | — |
| 200 | 0.86±.00 | 0.46±.04 | 0.04±.00 | 0.71±.00 | 0.73±.07 | 0.34±.01 | — | — | — |

### Extended Results

**Table 6.** Attacks against scDesign2 (no added noise).

| Method | 2 | 5 | 10 | 20 | 50 | 100 | 200 | 490 |
| --- | --- | --- | --- | --- | --- | --- | --- | --- |
| **OneK1k** |  |  |  |  |  |  |  |  |
| WB+aux | 1.00 | 0.99 | 0.96 | 0.94 | 0.85 | 0.78 | 0.64 | 0.64 |
| WB+aux (ClassB) | 1.00 | 0.95 | 0.89 | 0.95 | 0.85 | 0.76 | 0.73 | 0.68 |
| WB-aux | 1.00 | 0.93 | 0.84 | 0.81 | 0.73 | 0.59 | 0.58 | 0.52 |
| WB-aux (ClassB) | 0.30 | 0.42 | 0.47 | 0.46 | 0.55 | 0.51 | 0.52 | 0.50 |
| BB+aux | 1.00 | 0.81 | 0.79 | 0.70 | 0.66 | 0.57 | 0.59 | 0.55 |
| BB+aux (ClassB) | 1.00 | 0.97 | 0.89 | 0.89 | 0.79 | 0.72 | 0.65 | 0.59 |
| BB-aux | 1.00 | 0.68 | 0.66 | 0.64 | 0.63 | 0.54 | 0.59 | 0.51 |
| BB-aux (ClassB) | 0.25 | 0.39 | 0.46 | 0.46 | 0.55 | 0.51 | 0.57 | 0.51 |
| LOGAN | 0.85 | 0.51 | 0.59 | 0.57 | 0.47 | 0.53 | 0.49 | — |
| GAN-Leaks | 0.85 | 0.75 | 0.59 | 0.58 | 0.55 | 0.51 | 0.50 | — |
| DOMIAS+KDE | 0.30 | 0.60 | 0.55 | 0.52 | 0.50 | 0.48 | 0.50 | — |
| MC | 0.55 | 0.62 | 0.52 | 0.53 | 0.51 | 0.47 | 0.52 | — |
| **AIDA** |  |  |  |  |  |  |  |  |
| WB+aux | 1.00 | 0.92 | 1.00 | 1.00 | 1.00 | 0.98 | 0.87 | — |
| WB+aux (ClassB) | 1.00 | 0.97 | 0.98 | 0.94 | 0.95 | 0.88 | 0.82 | — |
| WB-aux | 1.00 | 0.92 | 1.00 | 0.96 | 0.81 | 0.68 | 0.63 | — |
| WB-aux (ClassB) | 0.55 | 0.67 | 0.70 | 0.58 | 0.54 | 0.51 | 0.54 | — |
| BB+aux | 1.00 | 0.92 | 1.00 | 1.00 | 0.96 | 0.92 | 0.77 | — |
| BB+aux (ClassB) | 1.00 | 0.96 | 0.95 | 0.90 | 0.90 | 0.82 | 0.77 | — |
| BB-aux | 1.00 | 0.92 | 0.97 | 0.87 | 0.69 | 0.60 | 0.60 | — |
| BB-aux (ClassB) | 0.55 | 0.66 | 0.69 | 0.56 | 0.53 | 0.51 | 0.53 | — |
| LOGAN | 0.50 | 0.58 | 0.37 | 0.52 | 0.54 | — | — | — |
| GAN-Leaks | 0.81 | 0.61 | 0.66 | 0.63 | 0.59 | — | — | — |
| DOMIAS+KDE | 0.44 | 0.49 | 0.51 | 0.49 | 0.52 | — | — | — |
| MC | 0.69 | 0.36 | 0.66 | 0.53 | 0.48 | — | — | — |
| **HFRA** |  |  |  |  |  |  |  |  |
| (11) |  |  |  |  |  |  |  |  |
| WB+aux | 0.95 | 0.94 | 0.84 | 0.84 | — | — | — | — |
| WB+aux (ClassB) | 0.70 | 0.59 | 0.76 | 0.72 | — | — | — | — |
| WB-aux | 0.85 | 0.74 | 0.65 | 0.65 | — | — | — | — |
| WB-aux (ClassB) | 0.60 | 0.52 | 0.56 | 0.62 | — | — | — | — |
| BB+aux | 0.95 | 0.94 | 0.79 | 0.77 | — | — | — | — |
| BB+aux (ClassB) | 0.70 | 0.59 | 0.75 | 0.71 | — | — | — | — |
| BB-aux | 0.85 | 0.75 | 0.64 | 0.63 | — | — | — | — |
| BB-aux (ClassB) | 0.60 | 0.52 | 0.55 | 0.61 | — | — | — | — |
| LOGAN | 0.45 | 0.57 | 0.45 | 0.46 | — | — | — | — |
| GAN-Leaks | 0.95 | 0.60 | 0.50 | 0.50 | — | — | — | — |
| DOMIAS+KDE | 0.65 | 0.52 | 0.55 | 0.50 | — | — | — | — |
| MC | 0.55 | 0.52 | 0.61 | 0.59 | — | — | — | — |

**Table 7.** Fidelity and privacy for synthetic data generated from training data with **490 donors** (OneK1K). LISI↑: real/synthetic cell mixing; ARI↑: cell-type structure; MMD*×*10^3^ ↓: distributional distance. BB*±*aux: scMAMA-MIA black-box without/with auxiliary data; *enhanced* variant with per-gene LLR evidence.

| SDG Method | DP Param | Fidelity |  |  | Privacy (scMAMA-MIA AUC) |  |  |  |
| --- | --- | --- | --- | --- | --- | --- | --- | --- |
| | | LISI↑ | ARI↑ | MMD$\times 10^3 \downarrow$ | BB-aux | BB+aux | *enhanced* BB+aux | |
| scDesign2 | (no DP) | 0.85 ± .00 | 0.43 ± .03 | 0.64 ± .42 | 0.51 ± .01 | 0.55 ± .02 | 0.59 ± .03 | |
| | $\eta = 10^{-4}$ | 0.00 ± .00 | 0.35 ± .03 | 312.56 ± 4.61 | 0.51 ± .03 | 0.51 ± .03 | 0.50 ± .03 | |
| | $\eta = 10^1$ | 0.00 ± .00 | 0.42 ± .02 | 308.68 ± 4.51 | 0.51 ± .03 | 0.52 ± .02 | 0.50 ± .03 | |
| scDesign3-V | | 0.34 ± .32 | 0.46 ± .02 | 36.92 ± 35.61 | 0.50 ± .01 | 0.55 ± .01 | 0.65 ± .04 | |
| scVI | | 0.90 ± .00 | 0.40 ± .04 | 0.26 ± .03 | 0.49 ± .01 | 0.63 ± .04 | 0.68 ± .03 | |
| ZINBWave | | 0.89 ± .00 | 0.40 ± .03 | 0.32 ± .07 | 0.50 ± .02 | 0.59 ± .02 | 0.61 ± .01 | |

**Table 8.** Fidelity metrics on the OK1K dataset for 10 and 20 donor settings. We report LISI (higher is better), ARI (higher is better), and MMD *×*10^3^ (lower is better) across synthetic data generation (SDG) methods and differential privacy (DP) noise levels. Results show consistent degradation in utility under stronger privacy regimes.

| Method | Setting | OK1K — 10 donors |  |  | OK1K — 20 donors |  |  |
| --- | --- | --- | --- | --- | --- | --- | --- |
| | | LISI↑ | ARI↑ | MMD$\times 10^3 \downarrow$ | LISI↑ | ARI↑ | MMD$\times 10^3 \downarrow$ |
| scDesign2 | no privacy | 0.882 | 0.496 | 0.427 | 0.879 | 0.500 | 0.407 |
| scDesign2 | $\eta = 10^{-4}$ | 0.885 | 0.484 | 0.419 | 0.881 | 0.456 | 0.399 |
| | $\eta = 10^{-3}$ | 0.882 | 0.474 | 0.404 | 0.880 | 0.504 | 0.396 |
| | $\eta = 10^{-2}$ | 0.835 | 0.524 | 0.422 | 0.842 | 0.510 | 0.366 |
| | $\eta = 10^{-1}$ | 0.738 | 0.570 | 0.543 | 0.726 | 0.525 | 0.454 |
| | $\eta = 10^0$ | 0.719 | 0.518 | 0.562 | 0.692 | 0.555 | 0.488 |
| | $\eta = 10^1$ | 0.714 | 0.539 | 0.555 | 0.682 | 0.532 | 0.496 |
| scDesign3-G |  | 0.901 | 0.487 | 0.277 | 0.891 | 0.464 | 0.252 |
| scDesign3-V |  | 0.896 | 0.483 | 0.313 | 0.892 | 0.477 | 0.290 |
| scVI |  | 0.913 | 0.439 | 0.347 | 0.907 | 0.434 | 0.318 |
| scDiffusion |  | 0.332 | 0.223 | 95.788 | 0.527 | 0.206 | 15.416 |
| NMF | no DP | 0.404 | 0.327 | 22.356 | 0.343 | 0.327 | 21.848 |
| NMF | $\epsilon = 10^3$ | 0.366 | 0.304 | 20.324 | 0.253 | 0.311 | 22.359 |
| | $\epsilon = 10^2$ | 0.370 | 0.295 | 20.498 | 0.245 | 0.311 | 22.622 |
| | $\epsilon = 10^1$ | 0.283 | 0.275 | 23.744 | 0.165 | 0.294 | 31.869 |
| | $\epsilon = 2.8$ | 0.019 | 0.293 | 66.479 | 0.035 | 0.290 | 56.870 |
| | $\epsilon = 10^0$ | 0.068 | 0.280 | 57.839 | 0.009 | 0.274 | 74.295 |

## Footnotes

1 https://bipress.boku.ac.at/camda2025/the-camda-contest-challenges/

2 typically a discriminative or generative model

3 scDesign2 applies a distributional transform to handle discrete count distributions [2].

4 but importantly, all generators for a trial are trained on the same *D*_*train*_.

5 LLMs aided in the organization of code, installing of external libraries, and running experiment suites. The core methods are owned and implemented by the authors.

6 https://cellxgene.cziscience.com/

7 Experiments are conducted on an AMD Ryzen Threadripper PRO 5965WX with 24 cores.

